# Intact RNA structurome reveals mRNA structure-mediated regulation of miRNA cleavage *in vivo*

**DOI:** 10.1101/2019.12.21.885699

**Authors:** Minglei Yang, Hugh C. Woolfenden, Yueying Zhang, Xiaofeng Fang, Qi Liu, Maria Louisa Vigh, Jitender Cheema, Xiaofei Yang, Matthew Norris, Sha Yu, Alberto Carbonell, Peter Brodersen, Jiawei Wang, Yiliang Ding

## Abstract

MicroRNA (miRNA)-mediated cleavage is involved in numerous essential cellular pathways. miRNAs recognize target RNAs via sequence complementarity. In addition to complementarity, *in vitro* and *in silico* studies have suggested that RNA structure may influence the accessibility of mRNAs to miRNA-Induced Silencing Complexes (miRISCs), thereby affecting RNA silencing. However, the regulatory mechanism of mRNA structure in miRNA cleavage remains elusive. Here, we investigated the role of *in vivo* RNA secondary structure in miRNA cleavage by developing the new CAP-STRUCTURE-seq method to capture the intact mRNA structurome in *Arabidopsis thaliana*. This approach revealed that miRNA target sites were not structurally accessible for miRISC binding prior to cleavage *in vivo*. Instead, the unfolding of the target site structure is the primary determinant for miRISC activity *in vivo*. Notably, we found that the single-strandedness of the two nucleotides immediately downstream of the target site, named Target Adjacent structure Motif (TAM), can promote miRNA cleavage but not miRNA binding, thus decoupling target site binding from cleavage. Our findings demonstrate that mRNA structure *in vivo* can regulate miRNA cleavage, providing evidence of mRNA structure-dependent regulation of biological processes.

## INTRODUCTION

MicroRNAs (MiRNAs) are ~21 nucleotide RNAs that impact various aspects of development and stress responses by post-transcriptionally regulating gene expression (1). MiRNAs are loaded onto ARGONAUTE proteins (AGO) to form functional post-transcriptional gene silencing effector complexes, miRNA-Induced Silencing Complexes (miRISCs) (2). miRISC is guided by the miRNA to target RNAs through sequence complementarity and cleave the target RNAs (3, 4). However, previous studies found that sequence complementarity is not the sole factor dictating miRNA cleavage (2). The structure of an RNA has been suggested to influence the silencing efficiency (5–7). However, these studies were unable to reveal native RNA structure features adopted through evolution because they indirectly assessed RNA structure by inserting a long sequence that was predicted to form a strong structure, such as a hairpin (5–7). Additionally, these structure assessments examined the target site together with its long flanking regions (5–7). This led to difficulties in dissecting the individual contributions from the different regions and confounded the identification of a specific RNA structure motif that regulated miRNA cleavage. Furthermore, these *in vitro* and *in silico* studies could not reflect the RNA structure folding status in living cells (8–10).

Recently, several transcriptome-wide structure probing methods for RNA *in vivo* have been established (8–10), which provide powerful tools to understand RNA structures under physiological conditions. However, these RNA structure methods that detect reverse transcription stalling (Structure-seq (9), icSHAPE (10, 11), Mod-seq (12)) or nucleotide mutation (DMS-MaP (13), SHAPE-MaP (14)) are not able to discern whether the chemical reactivity represents the RNA structure information for the endogenous degraded RNAs or for the intact RNAs (Supplementary Figure S1). Additionally, degraded mRNAs are capable of introducing false positive signals in the reverse transcription stalling methods because the 5’ end of the degraded mRNA will have an extremely high reverse transcription stalling signal (Supplementary Figure S1). Therefore, these methods are not able to reveal the causal relationship between RNA structure and miRNA cleavage.

To decipher the *in vivo* relationship between mRNA structure and miRNA cleavage, we have developed a novel method, CAP-STRUCTURE-seq, to obtain *in vivo* structures of target mRNAs before cleavage. We found that miRNA target sites were not structurally accessible *in vivo*. Instead, our analysis suggested that the unfolding of the target site structure is the primary determinant for miRISC binding prior to cleavage *in vivo*. Furthermore, by assessing the structure features flanking to the miRNA target sites, we also determined that the single-strandedness of the two nucleotides immediately downstream of the target site, which we named Target Adjacent structure Motif (TAM), can promote miRNA cleavage but not miRNA binding. Thus, TAM decouples target site binding from cleavage. Our study revealed the role of *in vivo* mRNA structure in the regulation of miRNA cleavage, providing evidence of mRNA structure-dependent regulation of biological processes.

## MATERIALS AND METHODS

### Plant materials and growth conditions

*A. thaliana* seeds of the Columbia (Col-0) and the *xrn4* mutant accession (15, 16) were sterilized with 70% (v/v) ethanol and plated on half-strength Murashige and Skoog medium (1/2 MS). The plates were wrapped in foil and stratified at 4°C for 3–4 days and then grown in a 22–24°C growth chamber for 5 days.

### Gel-based 18S rRNA structure probing

The gel-based method of structure probing used the same *in vivo* total RNA pools as for CAP-STRUCTURE-seq. To accomplish gel-based structure probing, reverse transcription was performed using 18S rRNA gene-specific DNA primers with 5’ end labelled Cy5 (TAGAATTACTACGGTTATCCGAGTA). The whole procedure was performed according to Ding, et.al(8). Each gel was detected by Typhoon FLA 9500 (GE Healthcare).

### (+)SHAPE and (−)SHAPE CAP-STRUCTURE-seq library construction

We modified the *in vivo* chemical probing protocol (8) by changing the reagent from dimethyl sulphate (DMS) to the SHAPE reagent, 2-methylnicotinic acid (NAI). NAI was prepared as reported previously (17). Briefly, five-day-old *A. thaliana* etiolated seedlings were suspended and completely covered in 20 ml 1X SHAPE reaction buffer (100mM KCl, 40mM HEPES (pH7.5) and 0.5mM MgCl_2_) in a 50 ml Falcon tube. NAI was added to a final concentration of 150mM and the tube swirled on a shaker (1,000rpm) for 15min at room temperature (22°C). This NAI concentration and reaction time had been optimized to allow NAI to penetrate plant cells and modify the RNA *in vivo* under single-hit kinetics conditions (Supplementary Figure S2A). After quenching the reaction with freshly prepared dithiothreitol (DTT), the seedlings were washed with deionized water and immediately frozen with liquid nitrogen and ground into powder. Total RNA was extracted using RNeasy Plant Mini Kit (Qiagen) according to the manufacturer’s instructions, followed by on-column DNaseI treatment in accordance with the manufacturer’s protocol. The control group was prepared using DMSO (labelled as (−)SHAPE), following the same procedure as described above.

To capture the structure information around the cleavage site of miRNA target genes, we adopted the feature of 5PSeq (18). The whole CAP-STRUCTURE-seq procedure is illustrated in Figure 1. In our method, the (+)SHAPE and (−)SHAPE RNA samples were treated with Terminator™ 5′-Phosphate-Dependent Exonuclease (TER51020, EPICENTRE co.), which processively digests RNA with 5′-monophosphate ends, thereby leaving mRNAs with 5’cap structures (Supplementary Figure S2B). Following the 5’cap enrichment, polyA+ selection was carried out using the PolyA purist Kit (AmbionTM) leaving intact (pre-cleaved) mRNAs with enriched 5’cap and 3’poly(A) tails. The resultant mRNAs were subjected to library construction following the STRUCTURE-seq procedure on Illumina HiSeq 4000 (BGI). The name of CAP-STRUCTURE-seq refers to 5’<u>**CAP**</u>-enriched and 3’ poly(A)-enriched RNA <u>**structure seq**</u>uencing.

**Figure 1.**
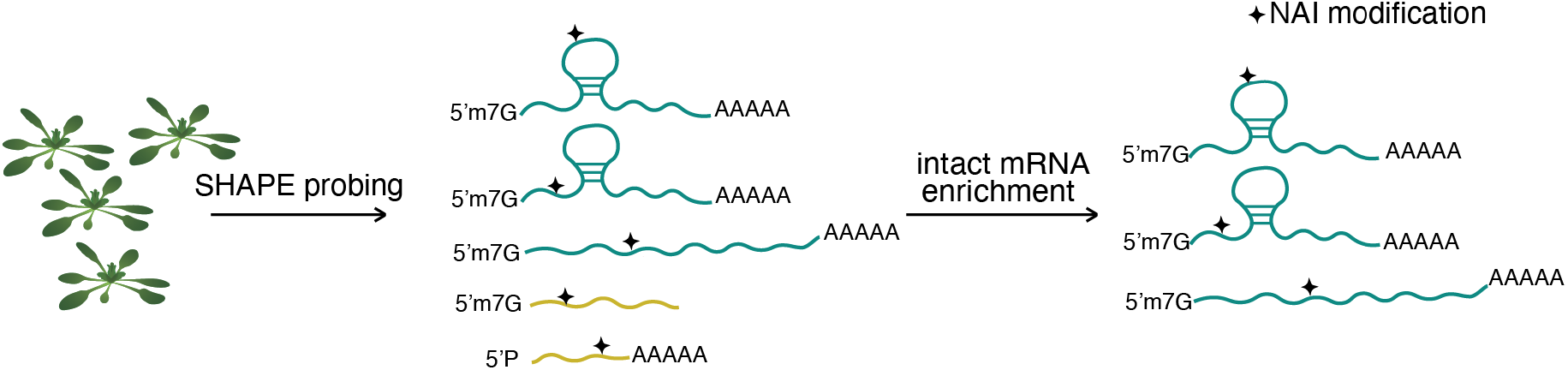
Schematic of CAP-STRUCTURE-seq. (+)SHAPE sequencing library generation showing NAI treatment, nucleotide modification and purification of intact mRNA steps. *A. thaliana* etiolated seedlings were treated with NAI. After extraction of total RNA, degraded mRNAs (dark yellow) were removed, leaving intact mRNAs characterized by 5’CAP and 3’ polyA+ (dark blue). cDNAs were obtained and subjected to an established library construction.

### CAP-STRUCTURE-seq analysis

We merged the biological replicates of the transcript-level reverse transcription (RT) stop counts to obtain a single (−)SHAPE library and a single (+)SHAPE library. We calculated the SHAPE reactivity using a slightly modified version of the formula in Ding et al.(8),

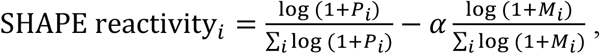

where *P*_*i*_ is the (+)SHAPE RT count and *M*_*i*_ is the (−)SHAPE RT count at nucleotide *i*. The factor, *α* (= min (1, Σ_*i*_ log(1 + *P*_*i*_) / Σ_*i*_ log (1 + *M*_*i*_)) is a simple library size correction factor. Setting *α*=1 recovers the reactivity formula in Ding *et al*.(8). The reactivities were then normalised using the box-plot method (19). For the SHAPE reactivity profiles, we extracted values in the 50 nucleotides upstream and downstream of target sites and calculated a per nucleotide mean and SEM.

### Degradome library construction

Five-day-old *A. thaliana* etiolated seedlings were grown as described above. They were collected and immediately frozen in liquid nitrogen and stored at −80°C. The seedlings were ground into powder. Total RNA was extracted using RNeasy Plant Mini Kit (Qiagen) according to the manufacturer’s instructions. On-column DNAaseI treatment was carried out according to RNase-Free DNase Set (Qiagen). To construct the Illumina library for degradome analysis, polyA+ selection was carried out using the Poly(A)Purist Kit (Ambion™). Selectively captured polyadenylated RNAs (1μg) were ligated directly to an DNA/RNA hybrid adapter (5’-CTACAC GACGCTrCrUrUrCrCrGrArUrCrUrNrNrN-3’) using T4 RNA ligase (NEB) at 37°C for 30 minutes. The ligated RNAs were subjected to RT by SuperScript III First-Strand Synthesis System (Invitrogen) with random hexamers fused with Illumina TruSeq adapters (5’-CAGACGTGTGCTCTTCCGATCTNNNNNN-3’). PCR amplification was performed on the ligated cDNA using Illumina TruSeq Primers. Two different barcode indices were used for two degradome biological replicates. The final dsDNA degradome libraries were subjected to next-generation sequencing on Illumina HiSeq 4000 (BGI).

### Degradome analysis

Raw reads were processed to remove 5’and 3’ adapter sequences. Degradome reads were mapped to the TAIR10 transcript reference and a degradome density file was generated. The degradation level of target genes was normalized by reads per kilobase per million mapped reads (RPKM).

### miRNA library construction

The same seedling samples stored at −80°C, as described above, were ground into powder using liquid nitrogen. Total RNA was extracted using mirVana miRNA Isolation Kit (Ambion™, Austin, TX, USA) following the manufacturer’s instructions. The integrity analysis was performed on a Bioanalyzer by the Beijing Genomics Institute (BGI), Shenzhen, China, which also performed the library construction according to standard protocols.

### miRNA-seq analysis

The small RNA sequences were processed by BGI to filter out the 5′ adapter sequences, 3′ adapter sequences and low-quality reads. We mapped two biological replicates against 253 miRNA sequences confidently annotated as *A. thaliana* mature miRNAs (20). We used Bowtie (21) for the mapping using the command ‘bowtie -f -a -S --best --strata -v 1’. pysam (21) was used to count the mapped reads.

### Cleavage efficiency (CE) calculation

The CE can be estimated by,

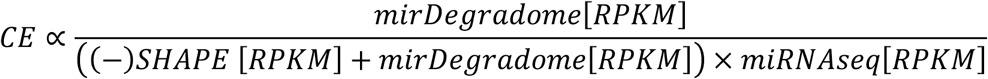

Briefly, we first identified how much miRNA expressed in our samples using the RPKM values from miRNA-seq. We used TargetFinder (22) to predict the miRNA target sites on the expressed transcripts in our samples and removed any duplicated target sites from the same miRNA family. TargetFinder predicts target sites with high specificity in *A. thaliana* by assigning a sequence complementarity penalty score (SCPS) (23) (Supplementary Figure S3A). Then, we mapped degradome reads to the reference transcripts that had been identified as miRNA target genes in *Arabidopsis* (20). We counted the reads within the target sites as the degradation products causing miRNA-mediated cleavage. Then, we summed the RPKM values from the (−)SHAPE and Degradome library to yield an estimate of how many transcripts served as substrates of miRNAs. The benefit of combining the (−)SHAPE and the degradome libraries to calculate the CE lies in its focus on miRNA-mediated cleavage events. The CE pipeline is illustrated in Supplementary Figure S3B and the derivation of the CE formula is described in the Supplementary Methods.

### Calculation of ΔG^⧧^_open_ and ΔG^⧧^_cutting_

ΔG^⧧^_open_ measures the energy required to open the target sites during miRISC binding. ΔG^⧧^_open_ was computed as the difference between the minimum free energy of the *in vivo* secondary structure and the minimum free energy of the ‘‘hard constrained’’ transcript, in which the target nucleotides were required to be unpaired (6, 24). By exploring a range of flanking region lengths upstream and downstream of the target site, we chose the upstream and downstream flank lengths to be 50 nucleotides for the majority of analyses. We used *RNAfold* from the Vienna RNA package (25) together with our SHAPE reactivity data to calculate the energy terms in ΔG^⧧^_open_, the RNA structures and the base pairing probabilities (BPP).

ΔG^⧧^_cutting_ measures the energy required to raise the initial substrate target RNA to the transition catalysis-compatible state, and is given by:

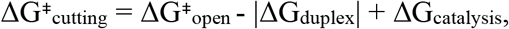

where ΔG_duplex_ is the binding free energy of the miRNA-target duplex, and ΔG_catalysis_ refers to the miRISC transition catalytic state energy. ΔG_duplex_ was calculated for the miRNA sequence and the target region sequence using *RNAduplex* from the Vienna RNA package (25).

The crystal structures of *Thermus thermophilus* Argonaute (TtAgo) (26, 27), human AGO2(28) and yeast *Kluyveromyces polysporus* Argonaute (KpAGO) (29) suggest that AGO proteins have a conserved catalytic mechanism. Furthermore, the transition cleavage model does not engage in any nucleotide interactions (26, 27). Therefore, we assumed that the activation energy, ΔG_catalysis_, is a constant for the same type of AGO protein. Quantum mechanics simulations estimate the value to be approximately 15 kcal mol^−1^(30). Therefore, ΔG^⧧^_cutting_ is given by:

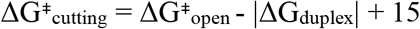

### Plasmid Construction

For cleavage efficiency validation, the miRNA156 target sites, followed by 0 or 2 Adenines (As) and ending with a G-quadruplex (GQS) or a stem-loop (SL) were synthesized and inserted into *Afl*II and *Pac*I of Firefly 3’UTR in vector inter2. We labelled the GQS constructs as 0A_GQS and 2A_GQS, and the stem-loop constructs as 0A_SL and 2A_SL, with the prefix indicating the number of Adenines. Antisense of miRNA156 target site constructs with the same flanking sequence were also synthesized as the control for each construct.

For the miRNA156 overexpression vector construction in AGO1 *in vivo* binding assay, the *MIR156B* genomic sequence was inserted into *Asc*I and *Sac*I of vector *pMDC32*. Primers are listed in Supplementary Table S1.

### Arabidopsis protoplast transformation

Protoplasts from the stable *MIR156* over-expression line were prepared and transformed according to the Tape-Arabidopsis Sandwich method (31). 16 h after transformation, protoplasts were centrifuged at 100 g for 2 min. RNA was extracted with Qiagen RNeasy kit and qRT-PCR quantification was performed with Bio-Rad CFX. Primers are listed in Supplementary Table S1.

### *In vivo* structure validation experiments

Four-week-old tobacco leaves were co-infiltrated with agrobacterium strains harboring plasmids of 0A_SL, 2A_SL, 0A_GQS or 2A_GQS. Two days after infiltration, the leaves were treated with 150 mM of the SHAPE reagent (NAI). The control group was treated with DMSO. Total RNA was extracted using RNeasy Plant Mini Kit (Qiagen) and then by on-column DNaseI treatment (following the respective manufacturer’s protocol). Gene-specific reverse transcription was performed as previously described by Kwok et al. (32), with a few modifications. 2 μg of *in vivo* total RNA was resuspended in 10 μl RNase-free water. Primer extension was performed with 2 pmol of DNA gene-specific primers (5’CATGCTTAACGTAATTCAACAGAAATTATATG) by Invitrogen SuperScript III reverse transcriptase. The resulting cDNA pellet was dissolved in RNase-free water and mixed with 1 μl 50 μm Mly1-HBLPCR-5’ssDNA linker modified by a 5’-phosphate and a 3’-3-Carbon spacer group (5’P-AGATCGACTCAGCGTCGTGTAGCTGAGTCGATCTNNNNNN-C3-3’), 10 μl Quick Ligase Reaction Buffer (2X), 1U Quick Ligase (New England Biolab) in a 20 μl system. The ligation was performed at 25°C for 1 h, followed by Phenol:Chloroform:Isoamyl Alcohol (25:24:1, v/v, sigma) and Chloroform:Isoamyl (24:1, v/v, sigma) purification.

The ligated cDNA samples were dissolved in 10 μl of water and used for the PCR reaction. The PCR reaction contained final concentrations of 0.5 mM VIC-labelled DNA gene-specific primers (the same as that used in the reverse transcription primers except the 5’ end was labelled with Vic), 0.5 mM of linker reverse primer (AGATCGACTCAGCTACACGACGC), 200 mM dNTPs, 1X ThermoPol reaction buffer and 1.25U of NEB Taq DNA polymerase in 25 μl. The solution was then extracted with Phenol:Chloroform:Isoamyl (25:24:1, v/v, sigma) and incubated with Mly1 restriction enzyme, according to the manufacturer’s protocol. Finally, the reaction pellets were dried and resuspended in Hi-Di formamide (Applied Biosystems/Life Technologies).

The Ned-labelled gene-specific primer (the same as that used in reverse transcription primer except the 5’ end was labelled with NED) was used to make sequencing ladders using linear DNA and 1 μl 5 mM ddTTP by Klenow DNA Polymerase I (New England Biolab) (33). Then, the reaction pellets were dried, resuspended in Hi-Di formamide (Applied Biosystems/Life Technologies) and run on an Applied Biosystems 3730xl Genetic Analyzer. The resulting data were analyzed using QuSHAPE (34).

### AGO1 *in vitro* cleavage assay

HA-tagged AGO1^WT^ was immuno-purified from *Arabidopsis* seedlings (35). The 0A_GQS and 2A_GQS designed RNAs were transcribed *in vitro* with T7 polymerase (NEB, 2040S) as substrates. To perform the slice assay, cleavage buffer (100mM ATP, 10mM GTP, 60mM MgCl_2_, 0.5M CPO_4_, 1mg/ml CPK) was added to 20μl beads in extraction buffer (1:1) bearing freshly purified HA-AGO1 from 3g seedling on the beads’ surface. 50 cps of labelled substrate was added to the reaction and incubated at 25℃. 10μl of the resultant liquid was added to 10μl 2x RNA loading buffer (95% Formamide, 0.02% SDS, 1mM EDTA, 0.02%, Bromothymol Blue, 0.01% Xylene Cyanol), denatured for 5min at 95℃ and loaded into a 1mm PAGE gel (10% acrylamides:bis 19:1, 7M Urea, 1xTBS). Then the gel was dried and exposed to a phosphor screen for image analysis.

### AGO1 *in vivo* binding assay

Four-week-old tobacco leaves were co-infiltrated with agrobacterium strains harbouring plasmids of *35S:MIR156B*, *35S:HA-AGO1*^DAH^ and 0A_GQS or 2A_GQS. Two days after infiltration, the leaves were collected and ground in liquid nitrogen. The protein/RNA complexes were extracted using two volumes of IP buffer (50 mM Tris-HCl pH 7.5, 150 mM NaCl, 5% β-mercaptoethanol, 1 mM EDTA, 10% glycerol, 0.1% NP-40, 1 mM PMSF, and 1X complete protease inhibitor cocktail). After removing insoluble debris by centrifugation, cell extracts were incubated with anti-HA antibody (Abcam ab9110) for 1h at 4℃ with gentle mixing. The anti-HA-decorated extracts were then incubated with pre-washed protein G magnetic beads for 1h. After incubation, the beads were washed 6 times with the IP buffer. The RNA produced after co-immunoprecipitation was precipitated with ethanol and glycogen, and analysed by RT-PCR. The miRNA156 expression levels were analysed by miRNA RT-PCR (36).

## RESULTS

### CAP-STRUCTURE-seq can accurately probe intact mRNA structures *in vivo*

To investigate how mRNA structure affects miRNA-mediated cleavage, RNA structure models should be captured before cleavage occurs. We therefore developed a novel strategy to obtain the structure of intact mRNAs, named CAP-STRUCTURE-seq (Figure 1). To obtain the RNA structure of intact mRNA, we performed *in vivo* selective 2′-hydroxyl acylation analyzed by primer extension (SHAPE) chemical probing on *Arabidopsis thaliana* (*A. thaliana*) with optimized conditions (Figure 1 and Supplementary Figure S2A). Next, we enriched the intact mRNAs through the terminator exonuclease treatment (Supplementary Figure S2B) (18), and then polyA+ purification to remove the degraded mRNAs. We generated two independent biological replicates of (+)SHAPE (samples with SHAPE treatment) and (−)SHAPE (control samples without SHAPE treatment) libraries according to the protocol described previously (8, 37). Between 90–97% of 340~380 million reads were mapped onto mRNAs (Supplementary Figure S4A, B) with the reproducibility of the CAP-STRUCTURE-seq library confirmed by comparing the two biological replicates (Supplementary Figure S5A, B). Nucleotide occurrence was consistent in both (−)SHAPE and (+)SHAPE libraries (Supplementary Figure S4C). To validate CAP-STRUCTURE-seq, we compared the SHAPE reactivity of the 18S rRNA with the corresponding phylogenetic covariance structure (Supplementary Figure S6A) and the 3D structure (Supplementary Figure S6B). We found that CAP-STRUCTURE-seq can accurately probe RNA structure *in vivo*, and in addition it outperforms the previous dimethyl sulphate (DMS)-based method, STRUCTURE-seq (8) (Supplementary Table S2).

To further validate CAP-STRUCTURE-seq we performed meta-property analyses with over 16,576 transcripts of sufficient RNA structure information (Supplementary Figure S7A). Our CAP-STRUCTURE-seq SHAPE reactivity data for *A. thaliana* exhibits similar genome-wide *in vivo* RNA structural properties to the previously results for a DMS-based method (8). For example, the region immediately upstream of the start codon showed particularly high SHAPE reactivity (Supplementary Figure S7B). This result further supports the notion that less structured regions near the start codon may facilitate translation (38, 39). Consistent with previous studies (8, 40), a periodic reactivity trend was found along CDS but was absent along UTRs (Supplementary Figure S7C). Similar to a RNase-based structure study in human (40), a unique asymmetric RNA structure signature at the exon–exon junction was also observed in *A. thaliana* (Supplementary Figure S7D). Taken together, these conserved RNA structure features suggest that CAP-STRUCTURE-seq successfully provides a global RNA structure model in plants.

We then assessed whether CAP-STRUCTURE-seq can overcome the limitations of previous transcriptome-wide RNA structure probing methods. The miRNA-mediated cleavage in the mRNA target site occurs at the tenth nucleotide of miRNA complementary sites (20), which leaves endogenous degraded products. In the previous DMS Structure-seq data, the cleavage site leads to reverse transcription stalling, and causes a skewed DMS reactivity profile due to false positive signals (Figure 2A). In our CAP-STRUCTURE-seq data, these degradation signals were excluded (Figure 2B), thereby overcoming the limitations of previous methods that include degradation products (Supplementary Figure S1) (8, 10, 13, 41, 42). Overall, these data demonstrate that CAP-STRUCTURE-seq can accurately identify *in vivo* structures of intact mRNAs.

**Figure 2.**
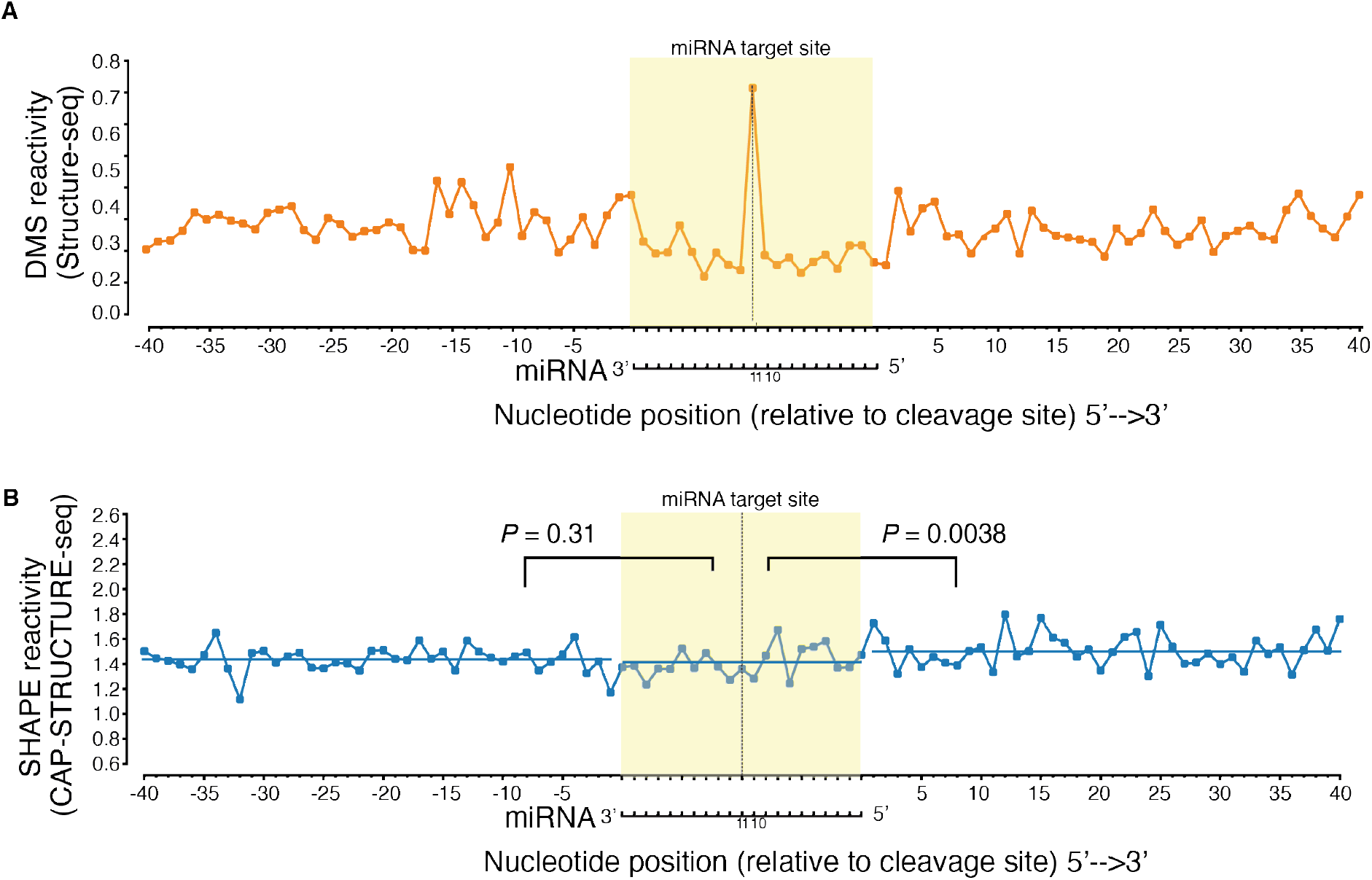
CAP-STRUCTURE-seq can accurately probe intact mRNA structures *in vivo*. **A**, Mean reactivity profiles across the miRNA target sites for the DMS reactivity from Structure-seq (8). The miRNA-mediated cleavage in the mRNA target site occurs at the tenth nucleotide of miRNA complementary sites, which leads to a skewed DMS reactivity profile at the eleventh nucleotide of miRNA complementary sites. **B,** Mean SHAPE reactivity profiles from CAP-STRUCTURE-seq. The miRNA cleaved products do not show a peak at the cleavage sites, indicating the elimination of false positive signals. In **A** and **B** the yellow shading indicates the target binding sites. The dotted lines refer to the 11^th^ nucleotide opposite to the miRNA. The horizontal axis is labelled from the 5’ end of the target gene to the 3’ end.

### The cleavage efficiency robustly measures miRNA-mediated cleavage events

Deciphering the *in vivo* relationship between mRNA structure and miRNA cleavage requires an *in vivo* structure model of target genes before cleavage, and the outcome after miRNA-mediated cleavage. Having developed a method to measure the former we turned our attention to the latter. To estimate the *in vivo* miRNA-mediated cleavage efficiency (CE), we drew inspiration from the definition of enzymatic activity (43). And we quantified CE by measuring how many degradation products were generated from one unit of substrate mRNA by one unit of miRNA (Methods, and Supplementary Methods). Our CE calculation is based on two underlying facts (20, 44): (i) miRNA-mediated cleavage is the major mRNA turnover pathway for target genes, (ii) the 5’ cleaved products are located within binding sites, which are transiently stable. Therefore, the degradation signal within target sites reflects the cleavage products from miRISC cleavage. We generated degradome libraries to estimate the degradation products (Supplementary Figure S3B and Methods) and miRNA-seq libraries to estimate miRNA abundance (Supplementary Figure S3B and Methods), with library reproducibility confirmed by comparing the biological replicates (Supplementary Figure S5C, D). We then combined the degradome, (−)SHAPE and miRNA-seq libraries to estimate CE (Supplementary Figure S3B, Methods and Supplementary Methods).

We verified the consistency of our CE against previously reported targets (Supplementary Table S3). For example, the CE of *AP2* targeted by miRNA172, which has been shown to act through translational repression rather than mRNA cleavage (45–47), was zero as expected (Supplementary Table S3). *SNZ* (Supplementary Table S3) is another target of miRNA172, and also showed no evidence of miRNA cleavage, consistent with the previous result (48). In contrast, *TOE2*, which is cleaved by miRNA172, had relatively high CE (Supplementary Table S3). Additionally, *TAS1a* and *TAS2*, which must be cleaved by miRNA173 to then serve as templates for trans-acting siRNA (tasi-RNA) (49), had high CE (Supplementary Table S3). These observations were consistent with their previous reported biological functions (45–49). Since sequence complementarity was reported to affect miRNA target cleavage (3, 4) we then systematically examined the relationship between sequence complementarity and CE. Globally, we found that sequence complementarity and CE were uncorrelated (Spearman correlation −0.015, Supplementary Figure S3E). In addition, targets with mismatches and/or GU wobble pairs (sequence complementarity penalty score, SCPS>0) were sometimes more effectively cleaved than targets with perfect complementarity (SCPS=0) (Supplementary Figure S3E). Our results indicate that other factors besides sequence complementarity between miRNA and mRNA may affect CE, with one possible example being mRNA structure. In summary, both the RNA structure of the intact mRNAs and miRNA cleavage *in vivo* can be quantitatively measured.

### The unfolding of the target site structure is the primary determinant for target RNA processing by AGO

With CAP-STRUCTURE-seq elucidating the RNA structure, we could begin to answer the elusive question about whether miRNA target sites were structurally accessible *in vivo*. Since our CAP-STRUCTURE-seq directly measures the *in vivo* structural accessibility via SHAPE reactivity (50), we assessed the SHAPE reactivity profiles across the miRNA target sites on the intact mRNAs. SHAPE reactivities of the target sites showed no significant difference from the upstream region (two-sided Mann-Whitney-U test, *P* value is 0.31, Figure 2B), and were lower than the downstream region (one-sided Mann-Whitney-U test, *P* value is 0.0038, Figure 2B). These features indicate that under physiological conditions the target sites are not highly accessible, which may provide a protective mechanism for target sites, mitigating against other cellular ribonucleases.

These relatively inaccessible target sites prompted us to ask whether the target site structure affects miRNA cleavage *in vivo*. To address this question, we examined two alternative energetic landscapes associated with the miRISC cleavage process *in vivo*: an enzyme-limiting scenario and a structure-limiting scenario (Figure 3A). In the enzyme-limiting scenario, the energy barrier (ΔG^⧧^_open_) between the inaccessible and accessible structural states (i.e., the unfolding of the target site) is lower than the barrier for catalytic cleavage (black line in Figure 3A). Thus, the target sites equilibrate quickly between inaccessible and accessible structural states during the binding step prior to the catalytic step of miRNA cleavage. In this scenario, the CE would vary with the free energy required to surmount the AGO catalytic barrier, ΔG^⧧^_cutting_ (Methods), and would be less affected by the RNA structure of the target site. In the structure-limiting scenario, the energy barrier (ΔG^⧧^_open_) between the inaccessible and accessible structural states is higher than the barrier for cleavage (red line in Figure 3A). Therefore, the target sites cannot achieve equilibrium binding with miRISC before catalytic cleavage. In this scenario, CE would vary with the free energy of opening the target site structure, ΔG^⧧^_open_, rather than ΔG^⧧^_cutting_. We used our *in vivo* structures to computationally approximate these two scenarios and explored a range of flanking lengths upstream and downstream of the target site (Figure 3B, C and Supplementary Figure S8A, B). Analysis of our SHAPE reactivity-informed structures revealed that, for most flank sizes, CE anti-correlated with ΔG^⧧^_open_ with a broad maximum centered around flanks of 50 nucleotides upstream and downstream (Spearman correlation of −0.23, *P* = 6.3e-9) (Figure 3B and Supplementary Figure S8A). However, for most flank sizes, CE had no correlation with ΔG^⧧^_cutting_ (Figure 3C and Supplementary Figure S8B), contrary to the reaction kinetics where the energy barrier is anti-correlated with reaction processivity. These results indicate that target site unfolding is the rate-limiting step that determines miRISC activity *in vivo*. Furthermore, this structure-limiting scenario reveals that the ribonuclease AGO undergoes “sticky regime” activation (51), where substrate mRNAs associate and dissociate with AGO more slowly than they are being cleaved.

**Figure 3.**
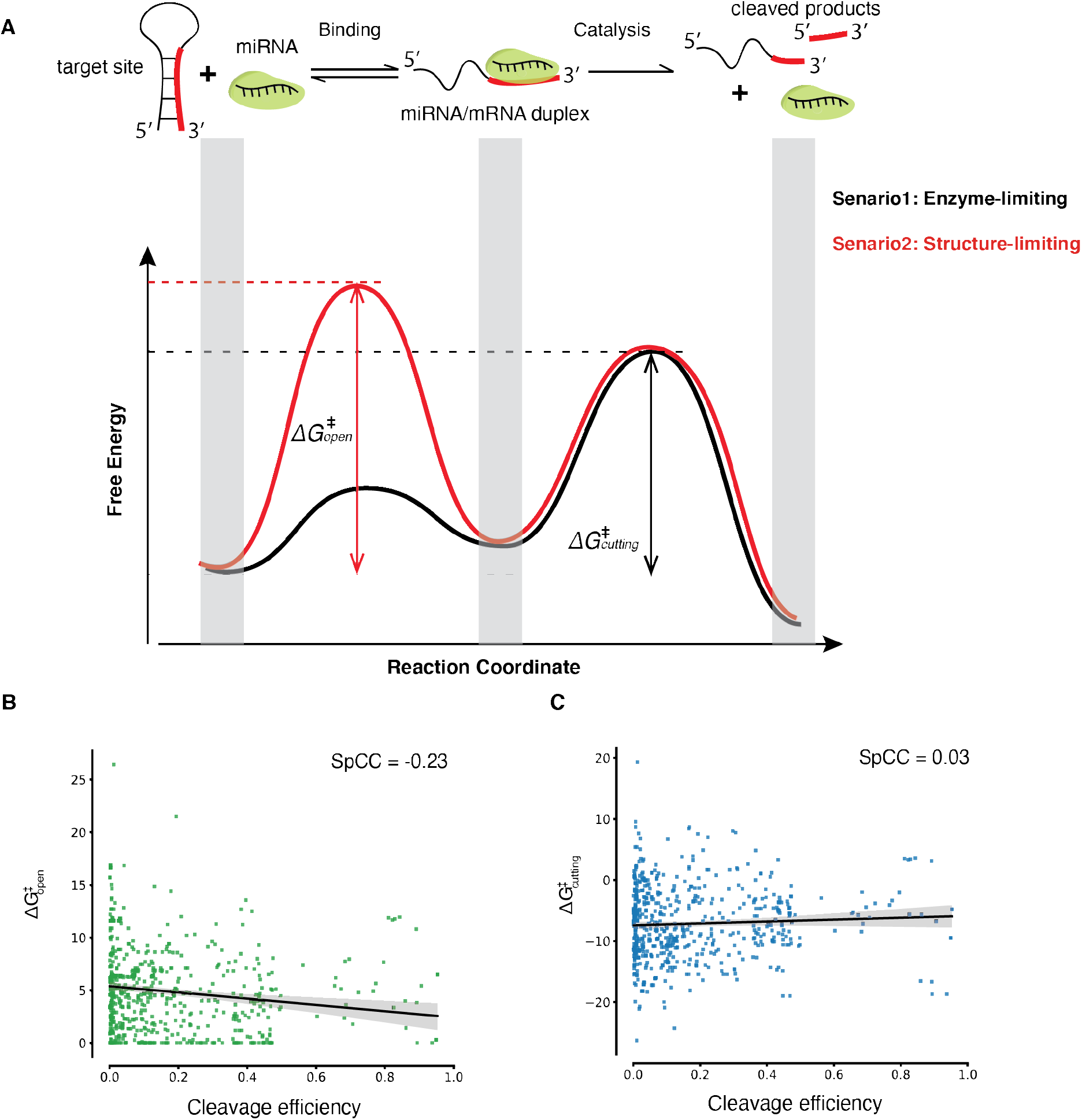
Energetic landscape of the miRISC cleavage process. **A,** The miRISC cleavage reactions include target binding and cleavage catalysis. Two alternative scenarios demonstrate the energetic landscape of miRNA cleavage (**black** and **red**). In the enzyme-limiting scenario (**black**), target site structure equilibrates quickly between inaccessible (closed) and accessible (open) states in the binding step compared to the catalytic step of miRNA cleavage. In this scenario, the apparent activation energy is ΔG^⧧^_cutting_, which measures the energy required to raise the initial substrate target RNA to the transition catalysis-compatible state. Alternatively, in the structure-limiting scenario (**red**), the target site cannot achieve equilibrium binding before cleavage. In this scenario, the energy barrier between the target site and the transient state is higher than the barrier for cleavage. And the apparent activation energy is equal to ΔG^⧧^_open_, which measures the energy required to open the target site structure. **B,** Spearman correlation between ΔG^⧧^_open_ and cleavage efficiency (647 target sites with the upstream and downstream flank lengths of 50 nucleotides, *P* = 6.3e-9 ***). **C,** A similar analysis to **B**, but for ΔG^⧧^_cutting_ and cleavage efficiency (*P* = 0.46).

The structure-limiting scenario implies that unwinding the miRNA target may be the limiting step in miRISC action, but once the miRNA is bound cleavage occurs. This is consistent with AGO RIP-seq results, where few target transcripts have been captured in wild type (WT) with less than 1 fold enrichment while target mRNA were enriched seven-fold in the catalytic mutant of (AGO1^DAH^) (52). We also found that ΔG^⧧^_open_ anti-correlated with the enrichment ratio of target RNAs from previous AGO1^DAH^-RIP-seq results (52) (Spearman correlation −0.21, *P* = 0.05). In contrast, the free energy of binding of the miRNA-target duplex (ΔG_duplex_) and ΔG^⧧^_cutting_ show no correlations with the enrichment (Spearman correlation 0.06 with *P* = 0.32 and −0.11 with *P* = 0.16, respectively). These observations suggest that the target sites are not structurally accessible *in vivo*, but rather the unfolding of the target site structure is the primary determinant for target RNA processing by AGO.

### Discovery of Target site Adjacent structure Motif (TAM) that contributes to miRNA-mediated cleavage *in vivo*

Having revealed that the target site structure affects cleavage *in vivo*, we then investigated whether the structure of the target site flanking regions is involved with miRNA cleavage. We assessed the RNA secondary structure by separating the RNA targets into non-cleaved (zero CE) and cleaved (positive CE) groups. We found higher SHAPE reactivity at the +1 and +2 nucleotides immediately downstream of target sites in the cleaved group relative to the non-cleaved group (Figure 4A), suggesting that these two nucleotides are more single-stranded than their neighbors. To confirm this observation, we used the SHAPE reactivity with the ViennaRNA *RNAfold* utility (53) to calculate the base pairing probabilities (BPPs). We found that the BPPs of the +1 and +2 nucleotides were much lower than their neighboring nucleotides (Figure 4B), indicating an increased likelihood of single-strandedness in the cleaved group compared to the non-cleaved group. Furthermore, the single-strandedness of the two nucleotides was unlikely to be due to sequence composition (Figure 5A) or AT content (Figure 5B) because there was no difference between the non-cleaved and cleaved groups. Our results reveal that a secondary structure feature, specifically single-strandedness of the two nucleotides adjacent to the 3’ end of the miRNA target site, generally exists *in vivo* in intact mRNAs that will undergo cleavage. We named this structure feature ‘Target Adjacent structure Motif’ (TAM).

**Figure 4.**
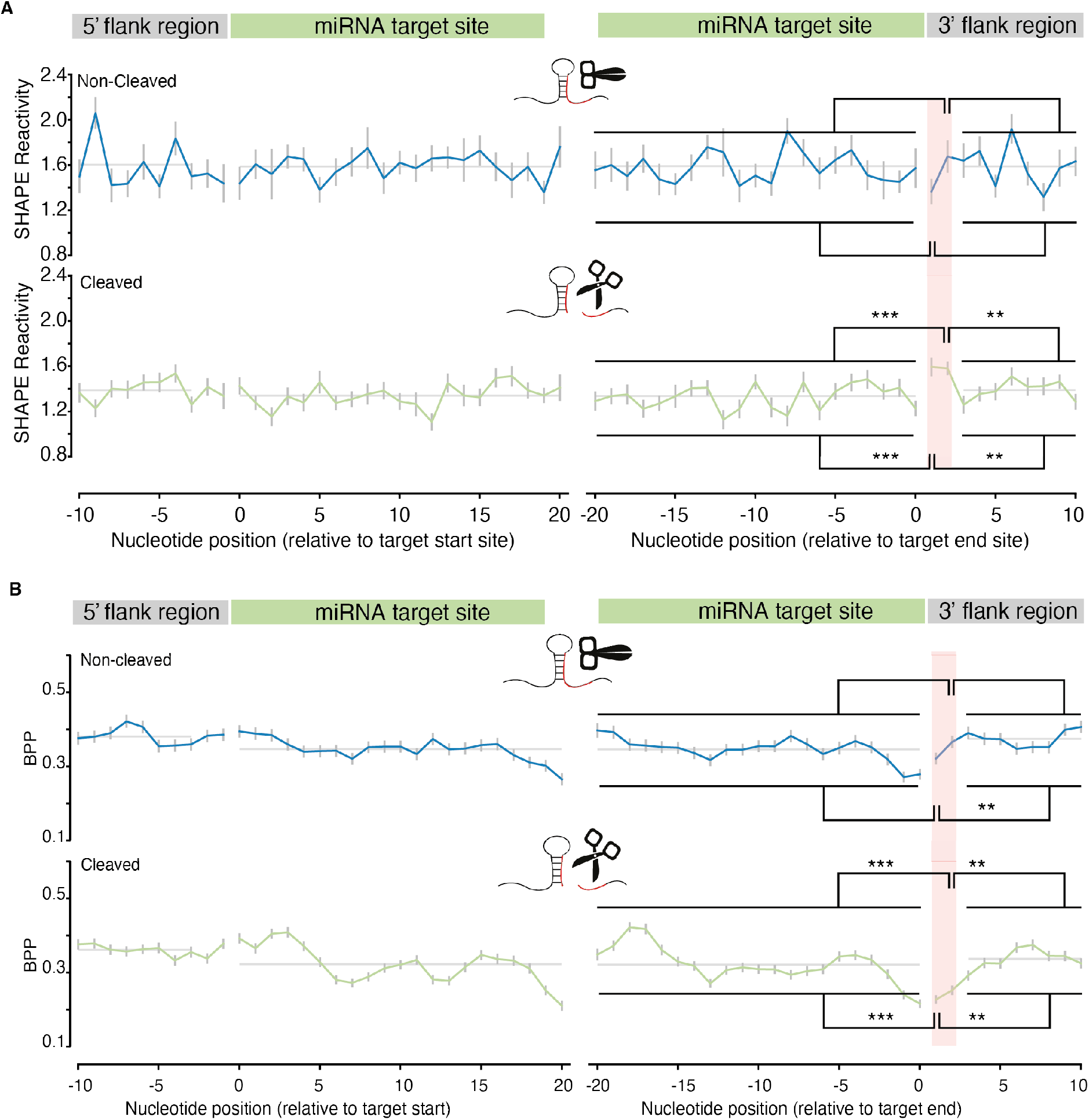
Discovery of a structure motif: <u>T</u>arget <u>A</u>djacent Structure <u>M</u>otif (TAM), which is independent of sequence composition. **A,** SHAPE reactivity profiles for miRNA target sites in the non-cleaved group (387 flanked target sites with reactivity values) and cleaved group (567 flanked target sites with reactivity values). The profiles show the per nucleotide mean +/− SEM across transcripts, aligned by target site start (left panels) and end position (right panels). Two nucleotides in the cleaved group (positive CE group), immediately downstream of target sites (TAM region), show significantly higher SHAPE reactivities compared to their neighbors (by Mann-Whitney-U tests). Compared to the upstream region (target sites), *P* = 4.8e-7*** for both 1^st^ and 2^nd^ nucleotides; Compared to the downstream region, *P* = 3.9e-3** for both 1^st^ and 2^nd^ nucleotides. The two individual nucleotides of the TAM region in the non-cleaved group (zero CE group) are not significantly higher than their neighbors by Mann-Whitney-U tests. B, Base pairing probability (BPP) at TAM in the non-cleaved group and cleaved group. SHAPE reactivity-directed base pairing probability (BPP) (Methods) for miRNA target sites in the non-cleaved group (387 target sites) and cleaved group (567 target sites). Corresponding to **A**, the two individual nucleotides within the TAM region (pink shading) show significantly lower BPP compared to the target site (*P* = 9.5e-7 *** for the 1^st^ nucleotide and *P* = 3.3e-6 ***) for the 2^nd^ nucleotide) and downstream region of TAM (*P* = 3.9e-3 **) for both 1^st^ nucleotide and 2^nd^ nucleotide) in the cleaved group. Overall, the two corresponding nucleotides in the non-cleaved group are not significantly lower than their neighbors by Mann-Whitney-U tests except the first nucleotide (*P* = 3.9e-3 **) compared to downstream regions). Since the SHAPE reactivity is the direct measurement of single-strandedness, here the subtle inconsistency between SHAPE reactivity and BPP may result from the uncertainty of the nearest neighbor parameter embedded in RNA structure prediction software (67). Comparison by Mann–Whitney U test.

**Figure 5.**
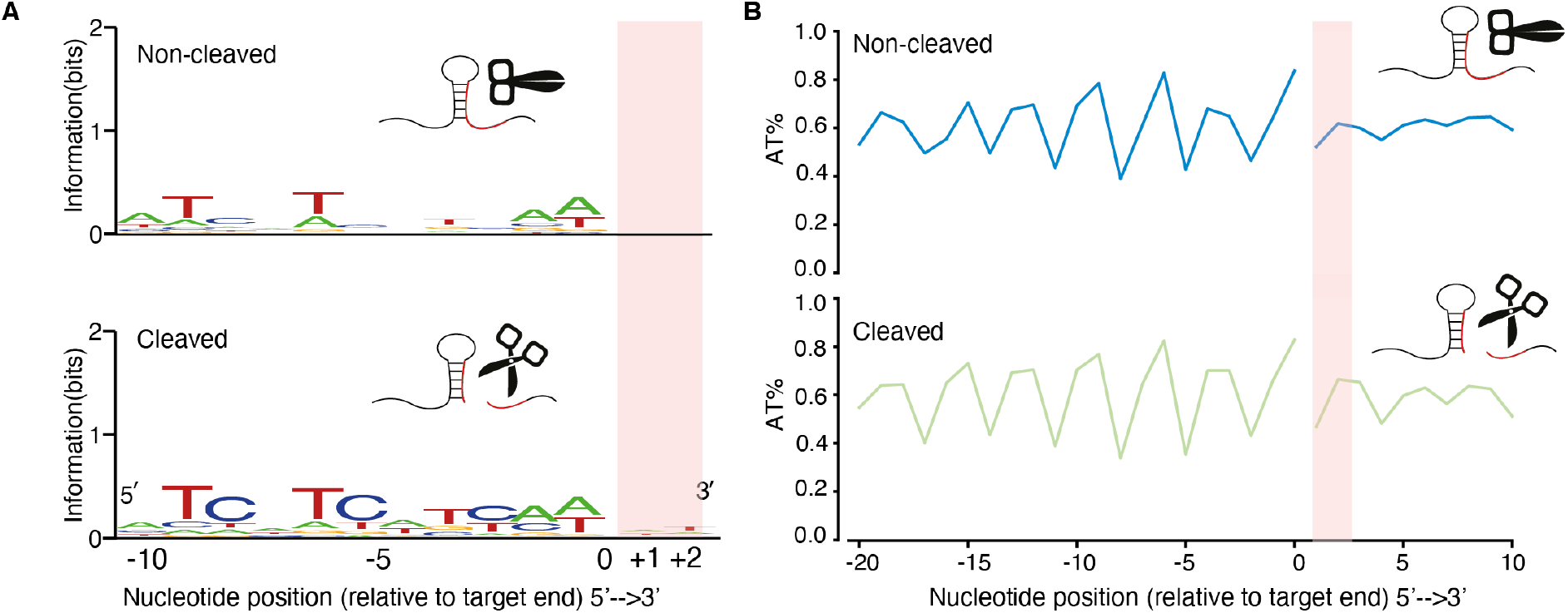
No sequence preference in the TAM region. **A,** Sequence composition around the target sites for the total cleaved (675 target sites) and non-cleaved groups (571 target sites). **B,** AT content around the target sites for the total cleaved (675 target sites) and non-cleaved (571 target sites) groups.

### TAM promotes miRNA cleavage but not miRNA binding

To explore the functional role of TAM in miRNA cleavage, we designed a structure assay (Methods) by concatenating miRNA156 target sites (20 nt) with a designed stable structure module (either a G-quadruplex structure or a stem-loop) to mimic the base-pairing state of the two nucleotides immediately downstream of the target site (Figure 6A and Supplementary Figure S9A). To maintain the single-strandedness of the TAM we inserted two Adenines (AA), or “slippery sequence”, between the target site and the designed stable structure module (Figure 6A and Supplementary Figure S9A). We confirmed the formation of TAM *in vivo* by using capillary electrophoresis (32) to resolve the *in vivo* RNA structure (Methods, Figure 6B and Supplementary Figure S9B). We then assessed the miRNA cleavage *in vivo* by measuring the levels of non-cleaved substrate mRNA. We found that the mRNA level of non-cleaved target genes with TAM was significantly lower (Student’s t-test *P* value < 0.01) than those without TAM (Figure 6C and Supplementary Figure S9C), which suggested increased cleavage when TAM was present.

**Figure 6.**
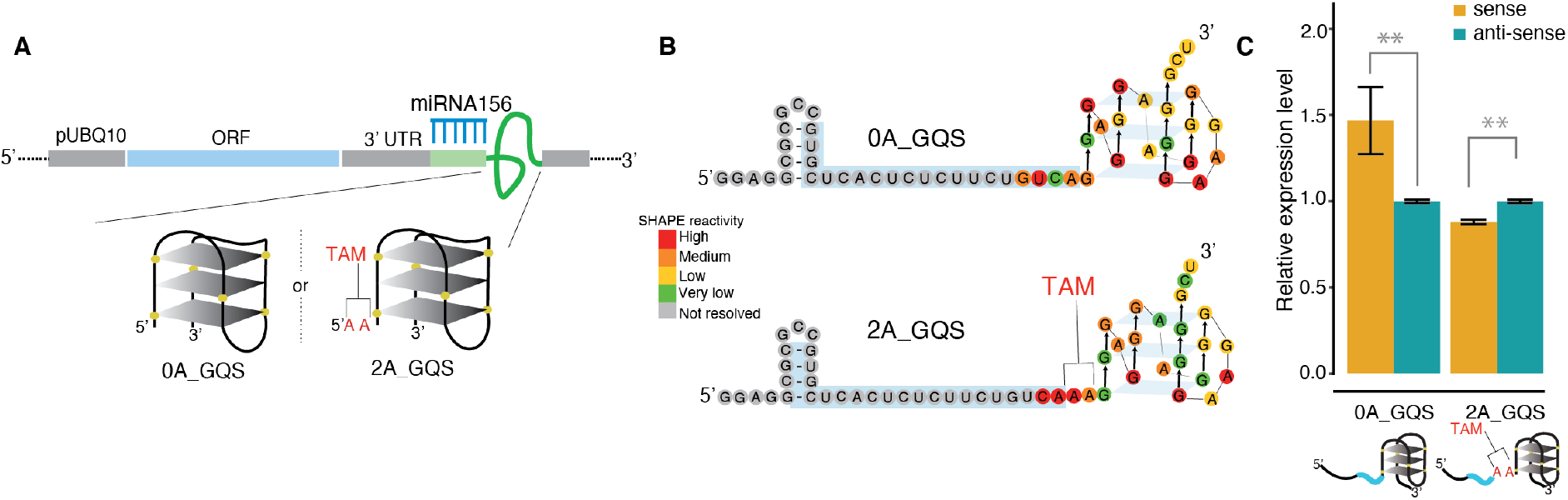
Validation of TAM functionality by a designed structure assay. **A,** Cartoon representation of the protoplast transformation assay to validate the TAM functionality using a designed structure assay. GQS refers to a G-quadruplex. The miRNA156 target sites (blue comb) followed by 0 or 2 Adenines (As) and ending with a GQS. The prefixes, “0A” and “2A”, indicate the number of Adenines. **B,** *In vivo* RNA structures of 0A_GQS and 2A_GQS. **C**, The non-cleaved mRNA abundance for the structures in **B** was measured by qRT-PCR (dark yellow bars). *P value* < 0.01 by Student’s t-test. The antisense target sites were used as controls (teal bars). Data are mean +/− SEM from three independent biological replicates.

To further confirm that the presence of TAM exclusively determines target cleavage, we performed an *in vitro* AGO cleavage assay using HA immuno-affinity-purified wildtype AGO protein and we found that the target RNA was cleaved only when TAM was present (Figure 7A). Our results reveal that TAM is essential for miRISC nuclease activity. One might expect TAM in the target mRNA to facilitate AGO binding instead of directly triggering the nuclease activity of AGO proteins. To test the possibility that TAM affects target binding, we conducted an *in vivo* binding assay (Methods) by using the slicing-defective AGO1 mutant, AGO1^D762A^. We found that AGO1 was able to bind the target RNAs with the same binding affinity irrespective of whether the TAM was present or absent (Figure 7B and C). Therefore, our data reveal that TAM regulates miRISC cleavage activity rather than affecting target binding.

**Figure 7.**
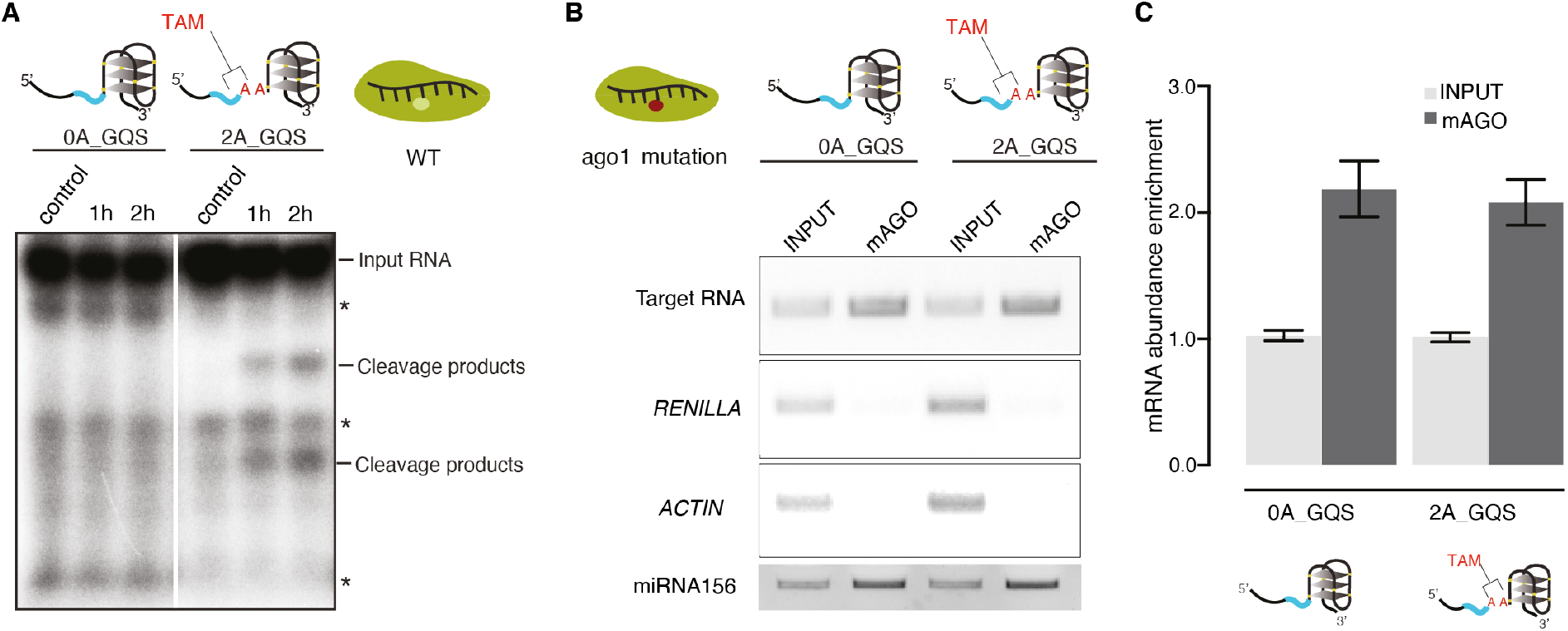
TAM promotes miRNA cleavage but not miRNA binding. **A,** *In vitro* AGO1 cleavage assay shows that target RNA is cleaved when the TAM is present. The target RNAs were incubated for 1h or 2h, where two cleavage products were present in the target with TAM on the X-ray film (as indicated). The asterisk indicates the background bands present in both control and experiment groups. **B,** *In vivo* AGO1 binding assay shows no difference between target RNAs with TAM and without TAM. *RENILLA* and *ACTIN* were used as the control. miRNA156 levels were measured in all the samples. **C,** The RNA abundance enrichment in **B** was quantified by amplicon intensities and normalized by input. Data are mean +/− SEM from three independent biological replicates.

## DISCUSSION

Here, we developed the novel CAP-STRUCTURE-seq method for differentiating RNA structure information for intact RNAs from degradation fragments. The intact RNA structurome facilitates the discovery of causal relationships between RNA structure and miRNA-mediated cleavage. In addition, we generated the first *in vivo* RNA structure landscape of *Arabidopsis* with structure information for all four nucleotides. The method outperforms the previous DMS-based Structure-seq method (8) (Supplementary Table S2) which only captured A and C structure information.

Target site accessibility has long been interpreted on the basis of spatial accessibility from a geometric viewpoint. However, in our study, we found that the target sites are not significantly spatially accessible *in vivo* (Figure 2B). Instead, we elucidated a structure-limiting scenario for miRNA cleavage (Figure 3A) from an energetic view. The spatially inaccessible target sites may provide a protective mechanism which prevents mRNAs from being targeted by other ribonucleases. Since miRISC has no helicase activity to unfold the RNA structure, miRISC has to take advantage of local structural variations, i.e., target site nucleotides become single-stranded (“breathing”), to find and bind its target site (Figure 3A). Thus, the equilibrium between a folded and an unfolded target site initially determines the binding rate (Figure 3A). This equilibrium is dependent on the energy cost of opening the target site (ΔG^⧧^_open_). Thus, the lower the energy barrier, the easier miRISC binds to the target sites (Figure 8).

**Figure 8.**
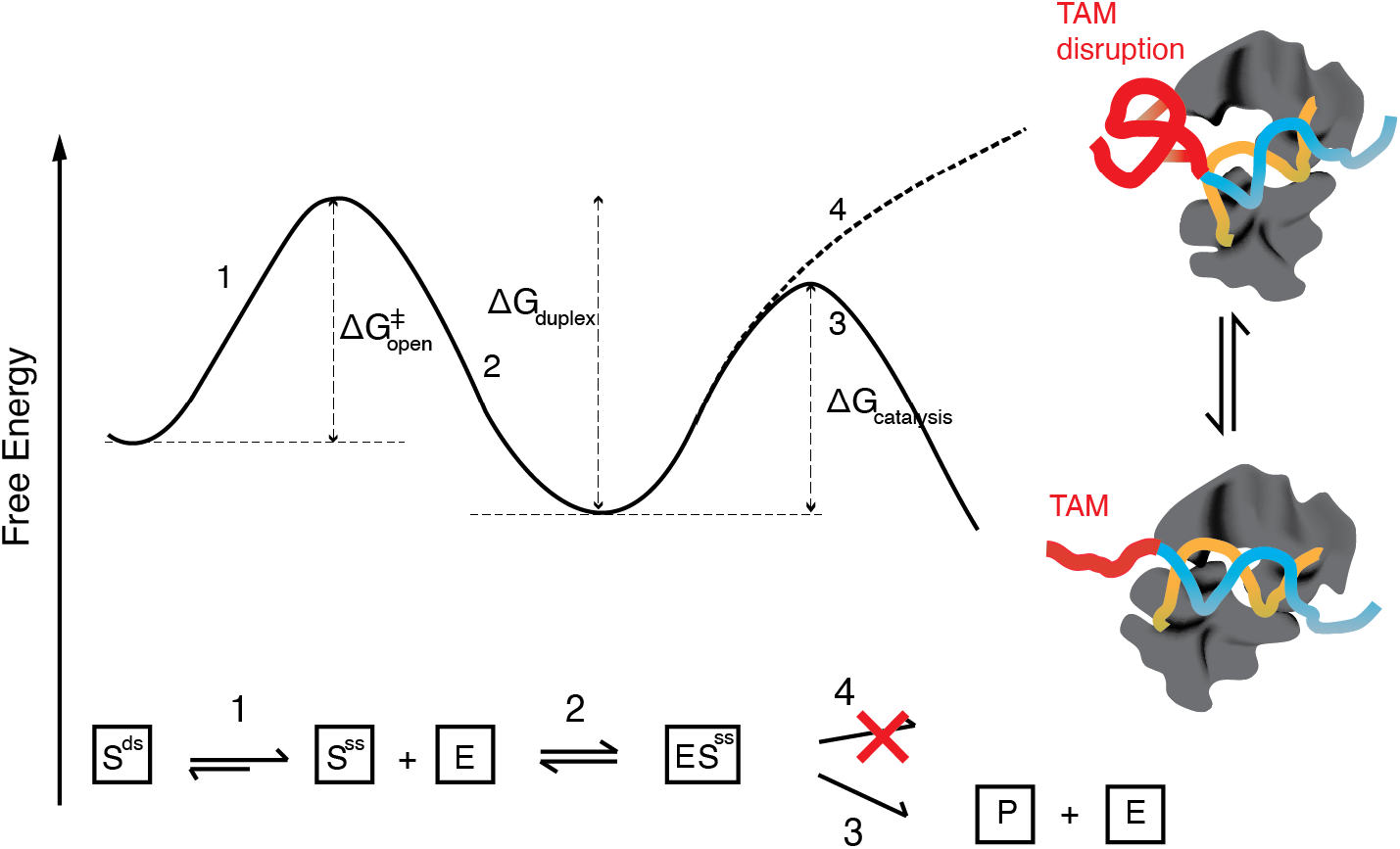
mRNA secondary structure-based model of miRNA-mediated cleavage. Illustration of the biochemical process of miRNA cleavage. (1) miRISC takes advantage of local structural variations, i.e., target site nucleotides become single-stranded (S^ss^,“breathing”) from double-stranded (S^ds^), to find and bind its target site; (2) miRISC (E) binds its target through base pairing (ES^ss^); (3) double-stranded TAM interacts with AGO protein and maintains the AGO in a cleavage-incompatible conformation; (4) the unwinding of the TAM can trigger the AGO into a cleavage compatible conformation, which then cleaves and releases the cleavage products (P). Energetic landscape of the RISC cleavage process corresponding to (A). (1) energy barrier for unwinding the target sites (ΔG^#^_open_); (2) energy released from miRNA-target duplex formation (ΔG_duplex_); (3) energy barrier for cleavage-compatible conformation with TAM (ΔG_catalysis_); (4) energy barrier for cleavage-incompatible conformation without TAM.

In living cells, many factors affect the final miRNA cleavage efficiency, including the miRNA precursor processing reviewed by (54), the miRNA methylation reviewed by (55), the miRNA exportation (56–58), miRNA localization and sequence complementarity reviewed by (59). Each factor contributes to the final miRNA cleavage efficiency. In our RNA structure study, we found that the target site unwinding (ΔG^⧧^_open_) can contribute 23% to the final miRNA-mediated cleavage *in vivo* (Figure 3B). Considering other contributors *in vivo*, our results suggested that under physiological condition the unfolding of the target site structure is the primary determinant for target RNA processing by AGO. In contrast, another factor, sequence complementarity does not show a high correlation with cleavage efficiency (Spearman correlation −0.015, Supplementary Figure S3E), indicating a relatively smaller contribution.

Once miRISC binds to the target sites, it needs to adjust the target conformation to perform the catalytic cleavage activity. A set of high resolution (~2.2 Å) ternary structures of *Thermus thermophilus* Argonaute (*TtAgo*) complexes have been solved (27), providing structural information about the transition between cleavage-incompatible and cleavage-compatible stages. AGO protein has been found to require conformation changes in three loops, L1, L2 and L3, to facilitate geometrical coordination of two magnesium ions (Mg^2+^) with the AGO nuclease activation site, the phosphate oxygens and in-line water, in order to facilitate the attack on the cleavable phosphate. The single-stranded TAM may promote this conformation transition and trigger the nuclease activity of AGO. Since the TAM is located at the 3’ end of the target site on the RNA, which is in parallel with 5’ end position of the miRNA, and the 5’ end of the miRNA interacts with the MID domain of AGO, we suspect that single-stranded TAM may engage or “touch” amino acids in the MID domain (Figure 8), thereby reducing the energy of conformation transition and facilitating the nuclease reaction (Figure 8).

The distinct roles of the target site and the TAM region decouples the target binding from target cleavage of miRISC *in vivo* (Figure 6, 7, 8 and Supplementary Figure S9). These properties are reminiscent of the CRISPR-CAS system (CAS9 and CAS13a) where both CAS9 and CAS13 decouple their binding and cleavage activity (60) (61) (62) (63). In addition, the endonucleolytic domains of CAS13 (HEPN domain), CAS9 (RuvC domain) (64) and RISC (PIWI domain) (65), contain an RnaseH-like fold and require Mg^2+^ as co-factor for catalytic activity. Furthermore, TAM can trigger the nuclease activity of miRISC. This mechanism, termed “substrate-dependent enzyme activation”, has also been found for CAS9 (61). This similarity indicates there may be a conserved mechanism between CAS system and miRISC system.

Our work indicates that the accessibility of the mRNA target site is the primary determinant for RISC endonuclease efficacy and that a structural motif within the pre-cleaved mRNA site is required to direct RISC cleavage. Adaptation of this motif within the *Arabidopsis* genome appears to have selected mRNAs that are readily cleavable, versus sites where miRNAs can bind, but miRISC cleavage does not occur. The presence of the TAM appears to be a prerequisite for RISC cleavage of its target mRNA and such knowledge has the potential to allow adjustment of the cleavability of RISC targets, potentially switching their mode of regulation. This supports the burgeoning hypothesis that RNAs may regulate RNA-binding protein (RBP) function rather than be regulated by RBPs (66). Furthermore, our results indicate that messenger RNA secondary structure has important physiological functions in many biological processes.

In summary, by deciphering intact mRNA structures *in vivo* through CAP-STRUCTURE-seq, we found that miRNA target sites were not structurally accessible *in vivo* and we demonstrated that the unfolding of the miRNA target site structure predominantly affected miRISC activity *in vivo*. Furthermore, we discovered that the native RNA structure motif, TAM, was sufficient to regulate miRNA cleavage *in vivo*. The mechanism that we found here provides evidence of mRNA structure-dependent regulation of biological processes *in vivo*. Our study reveals that *in vivo* mRNA structure serves as an additional regulator of miRISC activity, which will also facilitate the biotechnological engineering of gene silencing, and possibly provide an additional avenue towards crop improvement.

## SUPPLEMENTARY DATA

Supplementary Information are available at NAR online. Sequencing data are deposited in the Sequence Read Archive (SRA) on the NCBI website under the accession number SRR8444115.

## ACKNOWLEDGEMENT

We thank Dame Prof. Caroline Dean (John Innes Centre), Prof. Giles Oldroyd (SLCU, Cambridge), Dr. Desmond Bradley (John Innes Centre) and the group members in Ding lab for discussions with this work. We thank Mr. Mirko Ledda (UC. Davis) for discussions on data analysis. We thank Jim Carrington’s lab for providing us with the AGO construct for performing the binding assay.

## AUTHOR CONTRIBUTIONS

M.L.Y. and Y.L.D. conceived the study. M.L.Y., Y.Y.Z. and Y.L.D. designed the experiments with assistance from Y.S., A.C., P.B. and J.W.W. M.L.Y. designed the analyses. M.L.Y, Y.Y.Z., Q.L., M.L.V., X.F.Y. and X.F.F. performed the experiments with assistance from P.B., J.W.W. and Y.L.D. H.C.W., J.C. and M.N. performed the analyses. M.L.Y. wrote the paper with input from all authors. M.L.Y., H.C.W., Y.Y.Z., and X.F.F. contributed equally to this work. Correspondence should be addressed to Y.L.D.

